# Reinke crystals are immunoreactive for purine-synthesizing metabolic enzymes

**DOI:** 10.64898/2025.12.18.695235

**Authors:** John Woulfe, Trevor Flood, Sharlene Faulkes, David G. Munoz

**Author notes:** Corresponding author: John Woulfe, Dept. of Pathology, The Ottawa Hospital, Civic Campus, 1053 Carling Avenue, Ottawa, Ontario, Canada, K1Y 4E9.

## Abstract

Reinke crystals are a defining histological feature of human adult Leydig cells, the testosterone producing cells of the testis. These structures are present in the cytoplasm and the nucleus and display quantitative alterations in a variety of physiological and pathological contexts. The functional significance and protein composition of Reinke crystals have remained elusive for over a century. Here, we demonstrate that Reinke crystals are intensely immunoreactive for inosine monophosphate dehydrogenase (IMPDH), and phosphoribosyl pyrophosphate synthetase (PRPS), two key rate-limiting enzymes in the *de novo* synthesis of purine nucleotides. IMPDH and PRPS are two of several metabolic enzymes that are capable of forming mesoscale filamentous aggregates as a mechanism to regulate enzyme activity. IMPDH is also able to form crystals *in cellulo*. Our observations link Reinke crystal formation to purine nucleotide metabolism in Leydig cells. We discuss how this novel finding may relate to the unique dependence of Leydig cells on guanyl-based purine nucleotides for testosterone synthesis. The results of this study may have important implications for understanding metabolic contributions to male reproductive disorders as well as offering a novel diagnostic and theranostic tool applicable to Leydig cell neoplasms.

## Introduction

Leydig cells, the testosterone-producing cells of the testis, are essential for male reproductive health ^1^. Reinke crystals, first described by Frederich B. Reinke in 1896 ^2^, are a defining cytological feature of both native and neoplastic Leydig cells (for a review, see ^3^). The presence of Reinke crystals in neoplastic Leydig cells is of diagnostic utility. They are well-circumscribed, hyaline eosinophilic structures using conventional hematoxylin and eosin staining and are also demonstrable in Masson’s trichrome, Giemsa, and Gram preparations ^4-7^. Reinke crystals are 2-3 μm in size on average but can measure up to 20 μm ^8^. Ultrastructurally, their morphology is consistent with true crystals, showing “long-range order” in which 5 nm parallel filaments are arranged in a regular array throughout the entire structure. The protein composition and functional significance of Reinke crystals have remained enigmatic. Their abundance is dependent on Leydig cell developmental stage and is altered in a variety of testicular disorders (see ^3^). To date, candidate proteins that have been implicated as Reinke crystal constituents include nestin ^9^ and a 3β hydroxysteroid/calretinin complex ^7,10^. However, definitive confirmation of these is lacking ^3^.

Inosine monophosphate dehydrogenase (IMPDH) catalyzes the first committed step in the *de novo* synthesis of guanine-based (guanyl) purine nucleotides guanosine mono-, di-, and triphosphate (GMP, GDP and GTP). These, and especially GTP, are essential building blocks of RNA and DNA, are used as signalling molecules in several molecular pathways and are incorporated into co-enzymes. There are two isoforms of IMPDH that share 83% amino acid identity and have differential expression patterns ^11^. IMPDH1 is expressed constitutively in many tissues whereas IMPDH2 expression is also ubiquitous but is upregulated in proliferating tissues ^12-15^. IMPDH is one of a growing list of metabolic enzymes capable of regulated assembly to form long filaments as a mechanism to adjust enzyme availability and activity ^16-18^. Each filament consists of linearly stacked homo-octamers. Filament formation serves to resist feedback inhibition, thereby sustaining guanyl nucleotide synthesis under conditions of increased demand ^19^. Individual enzyme filaments coalesce to form microscopically visible cytoplasmic or intranuclear filament bundles called “cytoophidia” (Gk. For “cellular snakes”) ^18^. Cytoophidia formation correlates with increased enzyme activity and thus increased guanyl nucleotide synthesis. Accordingly, microscopically-detectable IMPDH filament assemblies serve as a marker for guanyl nucleotide-dependent cells ^16,17,20^. Importantly, IMPDH can also form large intracellular crystals ^21^. This spectrum suggests a temporally sequential series of assembly states in which IMPDH crystallization represents the ultimate stage in GTP-dependent cells. Intriguingly, *in cellulo* IMPDH crystals share morphological features with Reinke crystals. We have provided evidence that IMPDH crystallization occurs *in vivo*. Specifically, we described IMPDH filaments as well as crystals/paracrystals within the nuclei of neurons in the human brain ^22,23^. Interestingly, another metabolic enzyme, phosphoribosyl pyrophosphate synthase (PRPS) which catalyzes one of the first steps in *de novo* purine nucleotide synthesis ^23-26^ co-localizes with IMPDH in these neuronal structures ^23,27^. This is consistent with studies demonstrating PRPS filamentation/cytoophidia formation in cells, including neurons ^24^.

Relative to other endocrine cells, Leydig cells are uniquely dependent on GTP for testosterone synthesis ^28^. The latter is initiated in response to luteinizing hormone (LH) via a G-protein-coupled receptor (GPCR). Interestingly, GPCR signaling has been linked to purine metabolism by regulating the assembly of a purine-synthesizing multienzyme complex (metabolon) known as the “purinosome” ^29,30^. LH-receptor binding on Leydig cells results in recruitment of GTP to replace GDP on the G-protein Gsα. GTP-bound Gs activates its effector, adenylate cyclase resulting in the generation of cAMP. The latter stimulates cholesterol translocation to mitochondria where it is metabolized to pregnenolone which in turn is converted to testosterone by enzymes in the endoplasmic reticulum ^1^. Whereas all endocrine cells utilize GTP signaling to some extent, adult Leydig cells lack redundancy in this respect and are exclusively dependent on the LH-GTP-cAMP signaling axis ^28^. In contrast, fetal Leydig cells as well as testosterone-producing adrenal cells, (both of which notably lack Reinke crystals), can exploit alternate, non-GTP-dependent mechanisms ^28,31^. Moreover, LH receptor signaling in Leydig cells is characterized by maximal testosterone production even when only a tiny fraction (typically <1%) of available LH receptors are occupied by a ligand ^32-34^. This is critical for male reproductive health because it allows the testes to maintain stable testosterone levels even under low of fluctuating circulating LH ^35,36^. Mechanistically, this is attributable to massive signal amplification. There is evidence that this, in turn, is at least partly related to a stoichiometric imbalance in the ratio of G-proteins to LH receptors whereby a single occupied receptor can activate a large number of G-proteins, resulting in a 50-fold magnification of the LH binding reaction ^37^. This contrasts with catecholamine receptor signalling in which a closer numerical relationship between hormone receptors and G-proteins exists. It seems reasonable to surmise that this relative excess of G-proteins in Leydig cells would require a correspondingly robust supply of GTP. In this context, Leydig cells are functionally constrained by the availability of GTP, rendering them uniquely vulnerable to disruptions in purine nucleotide homeostasis.

Recognizing 1) the dependence of Leydig cells on guanyl nucleotide metabolism for their function, 2) evidence that intracellular IMPDH macromolecular assemblies characterize GTP-dependent cells and 3) the shared morphology of Reinke crystals, *in cellulo* IMPDH crystals, and the IMPDH/PRPS structures we have described in neurons, we were compelled to investigate whether Reinke crystals are immunoreactive for IMPDH and/or PRPS.

## Materials and Methods

The study was conducted in accordance with the ethical standards of The Ottawa Hospital. Tissue blocks of formalin-fixed, paraffin-embedded tissue from four cases of testicular Leydig cell tumours with adjacent non-neoplastic testis were obtained from the tissue archive in the Division of Anatomical Pathology of the Ottawa Hospital. These were resected from four male participants aged 28, 29, 70, and 71. Diagnoses in all cases were well-circumscribed Leydig cell tumours confined to the testicular parenchyma. The tumours measured from 1.3 to 1.6 cm. A representative block from each case was sectioned at a thickness of 5 μm, and sections were mounted onto coated slides. Immunohistochemical staining was performed on tissue sections using the Leica Bond™ system. Sections were deparaffinized and pre-treated using heat mediated antigen retrieval with an EDTA buffer (pH 9) for 20 minutes. The sections were then incubated in one of the following primary antibodies: rabbit polyclonal anti-IMPDH1 (1:400; Proteintech #22092-1 -AP, Lot #00018215; Rosemont IL USA), rabbit polyclonal anti-PRPS1 (1:200; ProteinTech #15549-1-AP), or mouse monoclonal anti-PRPS1/2/3 (1:200; Santa Cruz clone A11, sc-376440) for 30 minutes at room temperature and detected using an HRP conjugated compact polymer system. Slides were then stained using DAB as the chromogen, counterstained with hematoxylin, mounted and cover-slipped. As an experimental control for antibody specificity, the polyclonal primary antibodies were pre-absorbed using a 20-fold molar excess of the immunizing peptides (ProteinTech #Ag17473 for IMPDH and #Ag7907 for PRPS), and incubated overnight at 4 degrees with gentle agitation which reduced staining. For method specificity, omission of the primary antibodies resulted in a complete absence of staining. Double-immunofluorescence staining was performed by incubating sections overnight at 4 degrees in a cocktail containing the polyclonal IMPDH (1:100) and monoclonal PRPS (1:200) antibodies. Sections were washed with 1XTBST and then incubated with goat anti-mouse 594 (#A11005, Invitrogen) and donkey anti-rabbit 488 (#A21206) secondary antibodies for 2 h in the dark at room temperature. This was followed by incubation with a quencher (Vector TrueView Autofluorescence Quenching Kit #SP-8400, Vector Labs) to decrease autofluorescence. Sections were then washed, incubated with 5 ug/ml of DAPI (ThermoScientific #62248) and coverslipped. Microscopic imaging of was performed using Nikon Eclipse 80i light microscope. Photomicrographs were acquired using a Nikon Digital Sight Fi-1 camera and processed using NIS Elements imaging software version 4.60. For immunofluorescence, image acquisition was performed using a Zeiss 880 Confocal Microscope equipped with a Plan-Apo 63 × 1.4 numerical aperture. Processing of Airyscan 32-channel raw data was conducted using Zen Black 2.3 SP1 software.

## Results

In all 4 cases, cytoplasmic Reinke crystals were visualized in hematoxylin- and eosin (H and E)-stained sections in both tumour cells and native Leydig cells (Figure 1). Their frequency varied among cases. Reinke crystals in the cytoplasm appeared as well-circumscribed eosinophilic structures measuring from approximately 2 to 30 μm (Figure 1A). They were often rod-shaped or rectangular although various more irregular morphologies were common. Larger Reinke crystals, particularly the rod-shaped structures, appeared to “stretch” the plasma membrane of the host cell. In this limited series, we did not detect consistent differences in the size or morphology of cytoplasmic Reinke crystals between neoplastic and non-neoplastic Leydig cells in H and E-stained sections.

**Figure 1.**
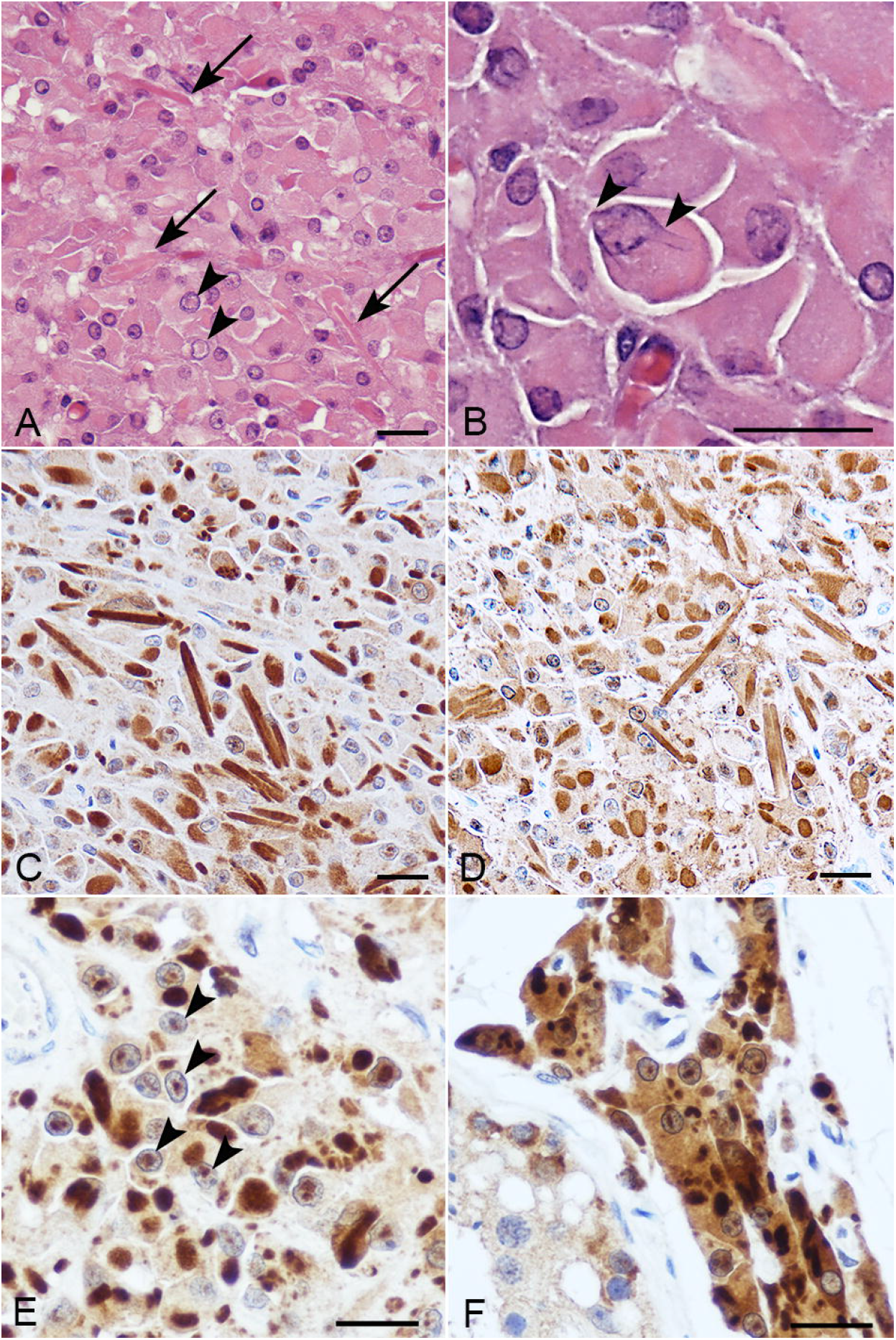
Reinke crystals are immunoreactive for IMPDH and PRPS. A-D: Reinke crystals in a Leydig cell tumour. A) Cytoplasmic (arrows) and nuclear (arrowheads) Reinke crystals are visible in an H and E stained section. B) A rod-shaped nuclear crystal extends to a length greater than the nuclear diameter, distorting and stretching the nuclear membrane (arrowheads). C, D) Immunostaining for IMPDH (C) and PRPS (D) reveals more abundant Reinke crystals morphologically identical to those visualized in H and E-stained sections. E) IMPDH immunohistochemistry is more sensitive for the demonstration of nuclear Reinke crystals. Many nuclear crystals (eg. arrowheads) stain with reduced intensity relative to cytoplasmic crystals. F) Both cytoplasmic and nuclear crystals are visualized in non-neoplastic Leydig cells using IMPDH immunohistochemistry. Scale bars=50 μm in A, C, and D; 25 μm in B, E, and F.

Reinke crystals were also visualized within the nuclei of neoplastic and non-neoplastic Leydig cells in H and E-stained sections (Figures 1A and B; arrowheads). The majority of these consisted of an amorphous eosinophilic area devoid of hematoxylin staining. In rare cases, nuclear crystals with a more solid rod-shaped morphology could be identified. Some of these were longer than the nuclear diameter and distorted the nuclear membrane (Figure 1B, arrow).

Across all four cases, both cytoplasmic and nuclear Reinke crystals displayed intense immunoreactivity for IMPDH and PRPS, concordant with the crystalline profiles seen on H&E (Figure 1B and C). All antibodies produced a similar intensity and distribution of immunostaining of cytoplasmic and nuclear Reinke crystals. In both Leydig cell tumours (Figure 1A-E) and native Leydig cell nests in the testicular interstitium (Figure 1F), Reinke crystals were far more evident and abundant in IMPDH and PRPS-immunostained sections relative to those stained using H and E (compare Figure 1A with C and D). Whereas less than half of the Leydig cell population appeared to have nuclear Reinke crystals in the latter, IMPDH or PRPS immunostaining revealed nuclear Reinke crystals in approximately 90% of the neoplastic Leydig cell population (Figure 1E). Moreover, immunostaining revealed features that were not appreciated on H and E staining. For example, whereas cytoplasmic crystals were intensely stained and showed well-defined edges, many nuclear crystals showed “blush” staining and ill-defined margins (Figure 1E, arrowheads). As shown in Figure 2, double-staining immunofluorescence confirmed that all cytoplasmic and nuclear Reinke crystals were immunoreactive for both IMPDH and PRPS.

**Figure 2.**
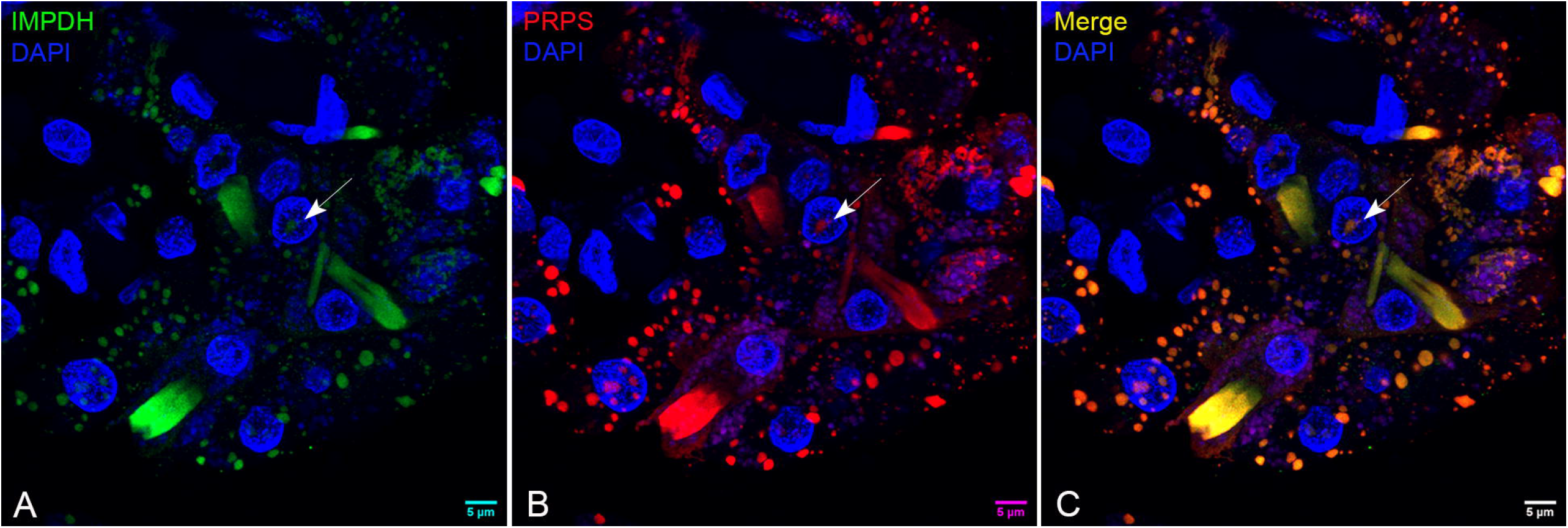
IMPDH and PRPS immunoreactivity are co-localized in Reinke crystals. Double immunofluorescence staining of neoplastic Leydig cells reveals immunoreactivity of cytoplasmic and nuclear (arrow) Reinke crystals for both IMPDH (A; green) and PRPS (B; red). Sections are counterstained with DAPI (blue). Scale bars=5 μm.

## Discussion

Despite having been described over a century ago ^2^, the protein composition of Reinke crystals has remained a mystery. Here, we demonstrate intense immunoreactivity of Reinke crystals for IMPDH and PRPS. These antibodies were generated using synthetic peptide immunogens corresponding to distinct but overlapping epitopes in each of the enzymes. Our study cannot distinguish the various IMPDH and PRPS isoforms. The target sequence of the polyclonal anti-IMPDH1 antibodies (amino acids 170-235) shares 77% sequence identity with the homologous domain of IMPDH2. Similarly, there are 3 highly homologous isoforms of PRPS (PRPS1, 2, and 3) in humans. The target epitope of the polyclonal antibody used in the present study (amino acids 1-318 of PRPS1) shares 98.7% and 98.1% sequence identity with PRPS2 and PRPS3, respectively whereas that of the monoclonal antibody (amino acids 143-270 of PRPS2) shares 98.7% and 96.5% sequence identity with PRPS1 and PRPS3, respectively. Interestingly, Keppeke and co-workers demonstrated co-localization of both IMPDH isoforms in cytoophidia, including intranuclear structures, in mammalian cells ^16^. Thus, it is possible that Reinke crystals contain multiple IMPDH and PRPS variants.

Among the novel findings of this study, we show that, like IMPDH, PRPS filaments are capable of incorporating into a crystalline array along with IMPDH. The co-assembly of distinct metabolic enzymes to form heterotypic assemblies is an emerging theme in the field of enzyme filamentation ^24,38^. Interestingly, co-assembling enzymes tend to be those located at critical “branch-points” in the purine biosynthetic pathway, including PRPS and IMPDH, suggesting that co-assembly may have an important regulatory role ^24^. In support of the possibility that these two enzymes represent the core component(s) of the filaments in the crystal lattice, in contrast to other proteins that have been nominated as Reinke crystal constituents, the ability of both IMPDH and PRPS to assemble into macrostructures, including, for IMPDH, crystals with a light microscopic and ultrastructural morphology similar to that described for Reinke crystals has been convincingly demonstrated. However, further studies, such as super-resolution microscopy, immunostaining studies at the electron microscopic level, and even cryoelectron microscopy are required to confirm this as well as to elucidate the spatial relationships between IMPDH and PRPS filaments within the crystals. ^21,39^.

By linking Reinke crystals to purine nucleotide metabolism in Leydig cells, our findings contextualize what was once merely a morphological curiosity within a coherent and physiologically relevant framework. As discussed above, Leydig cells are uniquely dependent on a sufficient supply of GTP. This is not only because they lack recourse to alternative non-guanyl-dependent signalling pathways ^28^, but also to satisfy the requirements of the signal amplification provided by the relative multitude of G-proteins which stoichiometrically outnumber their cognate LH receptors ^37^. Moreover, because Leydig cells respond to even small fluctuations in LH, GTP must be made available with high temporal fidelity. To meet these demands, we postulate that Reinke crystals provide readily accessible storage depots for IMPDH and PRPS to be rapidly deployed in response to sudden changes in GTP demand. This is analogous to the storage function ascribed to insulin crystals in the exocytotic vesicles of pancreatic beta cells and major basic protein 1 crystals in eosinophils ^40-42^. Consistent with this hypothesis, in conditions of higher LH levels and Leydig cell hyperplasia, including 17-βHSD deficiency and androgen insensitivity syndrome, Reinke crystals are reduced or not present ^3,43,44^. We propose that this reflects translocation of these key enzymes from the crystal to satisfy the requirements imposed by upregulated LH signaling in these scenarios. On the other hand, it is conceivable that Reinke crystals themselves provide a scaffold for guanyl-nucleotide synthesis in situ. This model would be predicated on the following: 1) that IMPDH and PRPS are enzymatically active in the crystalline context and 2) that the requisite enzymes catalyzing the upstream and downstream steps in the pathway are spatially proximal to the crystal. Although individual filaments contain enzymatically active oligomers, the status of IMPDH and PRPS functionality in the crystalline context is unknown. Future studies should explore the nucleotide-synthesizing capacity of Reinke crystals as well as their spatial relationship(s) with other purine-synthesizing metabolons, most notably, purinosomes ^30^. These hypothetical models are based on the premise that Reinke crystals are somehow involved in supplying sufficient GTP to meet the demands of Leydig cell signalling. The possibility that they are, alternatively or in addition, linked to some other unique, purine-related aspect of Leydig cell metabolism cannot be excluded.

Reinke crystals in the nucleus were also IMPDH and PRPS immunoreactive. Relative to H and E, immunohistochemical detection of these antigens revealed a much larger proportion of both neoplastic and native Leydig cells bearing nuclear inclusions. Nuclear Reinke crystals appeared to be less intensely stained and were more morphologically amorphous than the “rigid”-appearing cytoplasmic crystals. This may relate to observations suggesting that nuclear Reinke crystals may be at an earlier, precursor stage of formation, corresponding to paracrystalline structures ^4,45^. Whether these represent a nuclear “overflow” storage compartment of excessive cytoplasmic IMPDH/PRPS, or are functional is uncertain. In neurons, IMPDH/PRPS crystals show consistent localization adjacent to the nucleolus ^22^. The nucleolus consumes vast quantities of GTP for the synthesis of ribosomal RNA ^46,47^. It is tempting to speculate that juxtanucleolar crystals may provide a proximate “hyperlocal” reservoir of IMPDH/PRPS to be deployed for the synthesis of GTP. Alternatively, or in addition, IMPDH has been described in the cell nucleus where it has been ascribed a non-enzymatic “moonlighting” role ^48^. Specifically, nuclear IMPDH binds to DNA and functions as a transcription factor, regulating E2F-dependent gene expression in a cell-cycle dependent manner. In this context, Leydig cells might exploit the capacity of IMPDH to form crystals as a mechanism to dynamically regulate its nucleoplasmic availabilities to perform this moonlighting function.

The observation that children with Lesch-Nyhan syndrome, the prototypical disorder of purine metabolism, display impaired testicular development and testosterone deficiency ^49^ reinforces the link between Leydig cell function and purine metabolism in a pathophysiological context with potential implications for understanding the pathogenesis of male reproductive disorders. There may also be implications for cancer diagnosis and treatment. Leydig cell tumours are a rare form of testicular cancer. Most of these are benign although malignant variants can occur with metastatic potential. The identification of Reinke crystals in cytological preparations or tissue sections is useful in diagnosing these tumours ^6^. However, many tumours have very few crystals. In addition, Reinke crystals are sensitive to formalin fixation, contributing to false negative interpretation in H and E-stained sections ^7^. Immunohistochemistry for the detection of IMPDH or PRPS provides a novel, more sensitive histological marker for Reinke crystal identification in the diagnostic setting. With respect to therapeutic implications, several studies suggest that, under steady-state conditions, IMPDH filamentation is a hallmark of cells in which guanine nucleotide pools are limiting. This may represent a metabolic vulnerability that could be leveraged by IMPDH inhibitors to further deplete GTP, culminating in DNA replication stress and cell death. An important corollary of this is that the immunohistochemical detection of Reinke crystals in Leydig cell tumours may provide a histological biomarker predictive of a favourable therapeutic response to IMPDH inhibitors. Currently approved IMPDH inhibitors like mycophenolic acid as well as newer generation agents are receiving renewed attention as anti-neoplastic agents ^50^. Future preclinical studies and clinical trials should assess the susceptibility of Reinke-crystal bearing Leydig cell malignancies to IMPDH inhibition.

## Conclusions

Reinke crystals are immunoreactive for IMPDH and PRPS, linking these structures to purine nucleotide metabolism and possibly a role in regulating guanyl-nucleotide availability. In addition to solving a long-standing mystery, this finding has important implications for understanding the role of purine nucleotide metabolism in testicular function and disease, including cancer. Moreover, our previous demonstration of similar structures in neurons suggests that IMPDH/PRPS crystal formation may represent a paradigmatic organizational motif for understanding the regulation of purine nucleotide metabolism across diverse tissue types.

## Acknowledgements

The authors wish to acknowledge the technical support provided by the University of Ottawa Cell Biology Imaging and Acquisition Core (CBIA; RRID:SCR_021845) and the Louise Pelletier Histology Core Facility (RRID: SCR_021737), Department of Pathology and Laboratory Medicine, University of Ottawa. The authors specifically wish to acknowledge the excellent technical support provided by Liyuan Wang of the CBIA. No funding was received for conducting this study.

## Authors’ Contributions

J.W. designed the project, performed experiments, analyzed the data, and wrote the manuscript; T.F. provided tissue for the project, provided guidance regarding significance with respect to Leydig cell function and edited the manuscript; S.F. performed the immunohistochemical experiments and edited the manuscript; D.G.M. edited and provided critical input regarding organization of the manuscript. J.W. is the guarantor of this work and, as such, had full access to all of the data in the study and takes responsibility for the integrity of the data and the accuracy of the data analysis.

## Data Accessibility

All data pertaining to this study will be made readily available upon reasonable request.

